# Identification and Validation of a Previously Missed Mutational Signature in Colorectal Cancer

**DOI:** 10.1101/2025.06.06.658321

**Authors:** Mariya Kazachkova, Burçak Otlu, Marcos Díaz-Gay, Ammal Abbasi, Sarah Moody, Sandra Perdomo, David C. Wedge, Paul Brennan, Michael R. Stratton, Ludmil B. Alexandrov

## Abstract

Mutational signature analysis has greatly enhanced our understanding of the mutagenic processes found in cancer and normal tissues. As part of a recent study, we analyzed 802 treatment-naïve, microsatellite-stable colorectal cancers (CRC) and identified a *de novo* signature, SBS_D, which was conservatively decomposed into SBS18, a signature associated with reactive oxygen species. Here, we re-evaluate this decomposition and provide evidence that SBS_D represents a distinct mutational process from that of SBS18.

Through an independent analysis of 2,616 whole-genome sequenced microsatellite-stable CRCs across three distinct cohorts, we demonstrate that SBS_D is consistently present at a similar prevalence, suggesting that this signature may have been previously overlooked. Using a naïve decomposition approach, we demonstrate that the pattern of SBS_D better aligns with signatures previously associated with deficiencies in DNA polymerase delta (POLD1) proofreading and mismatch repair. However, multiple lines of evidence, including the absence of pathogenic mutations in the exonuclease domain of *POLD1* or in mismatch repair-associated genes, indicate that SBS_D is not driven by canonical defects in these DNA repair pathways.

Overall, this study identifies a previously unrecognized mutational signature in microsatellite-stable CRC and proposes that its etiology may be linked to DNA repair infidelity emerging late in tumor development in samples without canonical defects in DNA repair pathways.

## INTRODUCTION

Over the past decade, mutational signature analysis has become a standard component of genomics research^1,2^, greatly enhancing our understanding of the extrinsic and intrinsic mutagenic factors that influence the genomes of both cancer^3^ and normal somatic cells^4^. This has resulted in the development of global reference sets of mutational signatures, including those maintained by the Catalogue of Somatic Mutations in Cancer (COSMIC) database^5,6^.

In principle, mutational signature analysis can be performed in at least two ways: by assigning known reference signatures from databases like COSMIC to individual samples^7^ or by conducting *de novo* extraction of mutational signatures^8^. This latter approach is commonly employed in the analysis of large datasets containing at least several hundred samples, enabling the identification of novel mutational signatures by detecting *de novo* signatures that do not correspond to known references^8^. Specifically, all *de novo* extracted signatures are either directly matched to known reference signatures or, more frequently, decomposed into a combination of multiple reference signatures. Given the extensive catalog of known reference signatures, the decomposition process can be optimized to prioritize biologically relevant signatures, minimizing the risk of overfitting^9–11^. For example, a mutational signature associated with ultraviolet (UV) light exposure would typically be excluded from the decomposition of *de novo* signatures detected in colorectal or brain tumors as UV-induced mutations are not expected in these contexts^5^. Following decomposition, any signature that cannot be accurately reconstructed using a combination of known signatures with a cosine similarity of at least 0.90 is generally considered novel^9^.

In most cases, it is straightforward to determine whether a mutational signature is known or novel. Many *de novo* extracted signatures are easily decomposed into known reference signatures with high similarity. Conversely, some signatures exhibit patterns that cannot be fully reconstructed with substantial similarity, indicating the presence of a novel signature absent from the current reference set. Nevertheless, borderline cases exist where it remains uncertain whether a signature represents a truly distinct mutational process or merely a complex combination of previously known signatures. Scientifically, there is always a temptation to report a new signature, as it suggests the discovery of a potentially novel mutagenic process. However, validating such a discovery requires robust evidence in independent cohorts, particularly for signatures with unknown etiology, to ensure their validity. As a result, studies may adopt a conservative approach, potentially leading to the underreporting of genuinely novel mutational signatures.

Here, we revisit a recent *de novo* mutational signature analysis of 802 whole-genome sequenced microsatellite-stable colorectal cancers (CRC) from the Mutographs project^9^. In the original analysis, we identified a *de novo* single base substitution (SBS) signature, designated SBS_D, which had a cosine similarity of 0.90 when decomposed into known COSMICv3.4 reference signatures. At the time, we adopted a conservative approach and did not classify SBS_D as a novel signature^9^. Instead, we reported its decomposition into known COSMIC signatures, including SBS18, which has been associated with reactive oxygen species^12^. Here, we demonstrate that SBS_D represents a distinct mutational process from SBS18. Furthermore, we successfully extract SBS_D from three independent, previously published CRC cohorts^13–15^, none of which had identified or reported it. Finally, based on associations with known molecular phenotypes in CRC, we propose that SBS_D may result from POLD1 proofreading and mismatch repair infidelity occurring late in the evolution of tumors without detectable canonical defects in either pathway. Collectively, our findings reveal the existence of a previously unrecognized mutational signature in microsatellite-stable CRC samples and underscore the challenges associated with identifying novel mutational signatures.

## RESULTS

### Mutational Signatures and Their Predefined Schemas

All analyses of mutational signatures intrinsically rely on a predefined schema for classifying somatic mutations^12^. For somatic single base substitutions, the SBS-96 context is the most widely used classification schema and serves as the foundation for the current set of COSMIC reference SBS signatures^5^. In the SBS-96 framework, somatic mutations are classified into six base substitution types (C>A, C>G, C>T, T>A, T>C, and T>G) based on the pyrimidine base change within the mutated Watson-Crick base pair (*e.g.*, C>T representing a C:G>T:A mutation). Each of the six types is further subdivided into 16 subtypes by incorporating the trinucleotide context, which accounts for the adjacent 5′ and 3′ bases of the mutated pyrimidine. While SBS-96 is the most used schema for SBSs, higher-resolution schemas like SBS-288 and SBS-1536 have been applied in recent large-scale analyses, including in previous Mutographs studies^9–11,16^. These schemas enhance resolution by incorporating additional features, such as transcriptional strand information in SBS-288 or an extended pentanucleotide context in SBS-1536^17^ (**Methods**).

### Original Mutographs *De Novo* Signature Analysis

In the original analysis of 802 treatment-naïve, microsatellite-stable, DNA repair-proficient CRC samples from Mutographs, *de novo* extraction was performed using the SBS-288 classification schema^9^. This resulted in the identification of 16 *de novo* mutational signatures, labeled SBS_A through SBS_P (note that the use of letters distinguishes these from COSMIC reference signatures, which typically use numbers, *e.g.*, SBS18). Of these, 12 signatures were decomposed into COSMIC reference signatures, with the decomposition optimized to exclude signatures associated with defective DNA repair, as the cohort included only DNA repair-proficient samples. The remaining four signatures represented mutational patterns not previously cataloged in COSMICv3.4 and were, as such, classified as novel. One of the 12 *de novo* signatures that was decomposed into COSMIC signatures, SBS_D, achieved a borderline reconstruction cosine similarity of 0.90. However, there was insufficient evidence to classify it as a novel signature, as no other large CRC mutational signature study had previously reported it. Adopting a conservative approach and given insufficient evidence at the time, we reported SBS_D as a mixture of previously known signatures while acknowledging the possibility that it may represent a novel signature^9^. Importantly, SBS_D was found in 428/802 microsatellite-stable CRC samples, contributing 8.5% of all mutations (**Figure 1*a***).

**Figure 1:**
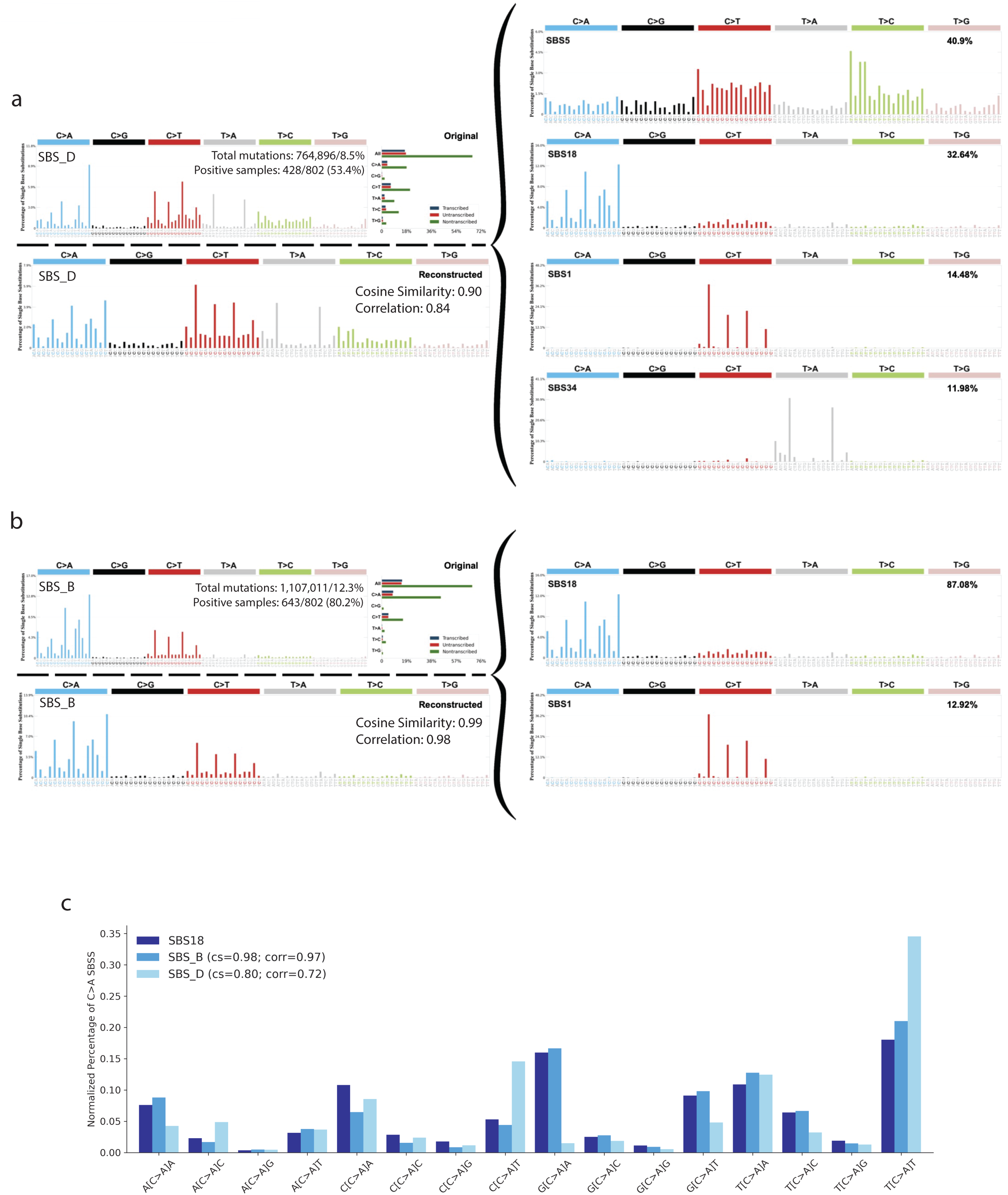
SBS_D and SBS_B from the Mutographs colorectal cancer cohort. ***(a)*** Optimized decomposition of SBS_D showing the total number of mutations attributed to the signature and the proportion of samples in which it is detected. ***(b)*** Optimized decomposition of SBS_B showing the total number of mutations attributed to the signature and the proportion of samples in which it is detected. ***(c)*** Comparison of the C>A contexts among COSMIC signature SBS18, *de novo* signature SBS_B, and *de novo* signature SBS_D. SBS_B and SBS_D were collapsed to the SBS-96 mutation classification schema, and normalization was performed within the C>A contexts. Cosine similarity (cs) and Spearman correlation (corr) are shown relative to the C>A contexts for SBS18.

### Reevaluating the Pattern of SBS_D

The pattern of SBS_D was decomposed into COSMICv3.4 signatures SBS1 (14.5%), SBS5 (40.9%), SBS18 (32.6%), and SBS34 (12.0%) (**Figure 1*a***). SBS1 and SBS5 are ubiquitous clock-like mutational signatures present in all types of cancer^12^ and normal tissues^18^. Due to their pervasive nature, some degree of SBS1 and SBS5 contamination is expected in the decomposition of any *de novo* signature^8^, and, as anticipated, these signatures were found in many *de novo* signatures extracted from the Mutographs CRC cohort^9^. As such, if the decomposition was accurate, SBS_D would represent a combination of the reactive oxygen species-associated signature SBS18, the unknown etiology signature SBS34, and contributions from ubiquitous clock-like signatures (SBS1 and SBS5), with SBS18 being the predominant non-clock-like contributor. However, another *de novo* signature extracted from this cohort, SBS_B, exhibited a near perfect match to SBS18 (**Figure 1*b***). Moreover, the C>A component, which constitutes the majority of SBS18 mutations, showed substantial differences between SBS18 and SBS_D (cosine similarity [CS]=0.80; Spearman correlation =0.72) but was nearly identical between SBS18 and SBS_B (CS=0.98; CORR=0.97; **Figure 1*c***). Considering the extremely high similarity between SBS18 and SBS_B, we will henceforth refer to SBS_B as SBS18^. Additionally, as part of the Mutographs study, additional extractions were performed using the SBS-96 and SBS-1536 classification schemas. Both approaches yielded two signatures closely resembling SBS18^ and SBS_D (**Supplementary** Figure 1). This consistency across different mutational schemas highlights the robustness of the patterns for these signatures, irrespective of the underlying classification framework. Overall, these findings raised doubts about the assignment of SBS18 as a contributor to SBS_D and suggested that SBS_D could represent a distinct mutational process.

### SBS_D Represents a Mutational Process Distinct from SBS18

To enhance our understanding of SBS_D and its relationship with SBS18^, we performed molecular timing on the microsatellite-stable, DNA repair proficient Mutographs CRC samples where copy number information was available (774 of 802 samples; 96.5%) by categorizing SBS mutations based on their timing relative to copy number gains during cancer development into two groups: early clonal and late clonal (**Methods**). This stratification allowed us to analyze the contributions of SBS_D and SBS18^ across distinct stages of tumor evolution. Our analysis revealed that SBS_D was preferentially active in late clonal mutations compared to early clonal mutations (p-value<0.0001; **Figure 2*a***). In contrast, SBS18^ was typically active exclusively early or both early and late but was rarely observed exclusively in late clonal mutations (p-value<0.0001, **Figure 2*b***). Notably, prior studies have similarly demonstrated that SBS18 is either preferentially early or constant in CRC development^19,20^. These findings indicate that SBS_D is an evolutionarily late signature, likely representing a mutational process distinct from that of SBS18^.

**Figure 2:**
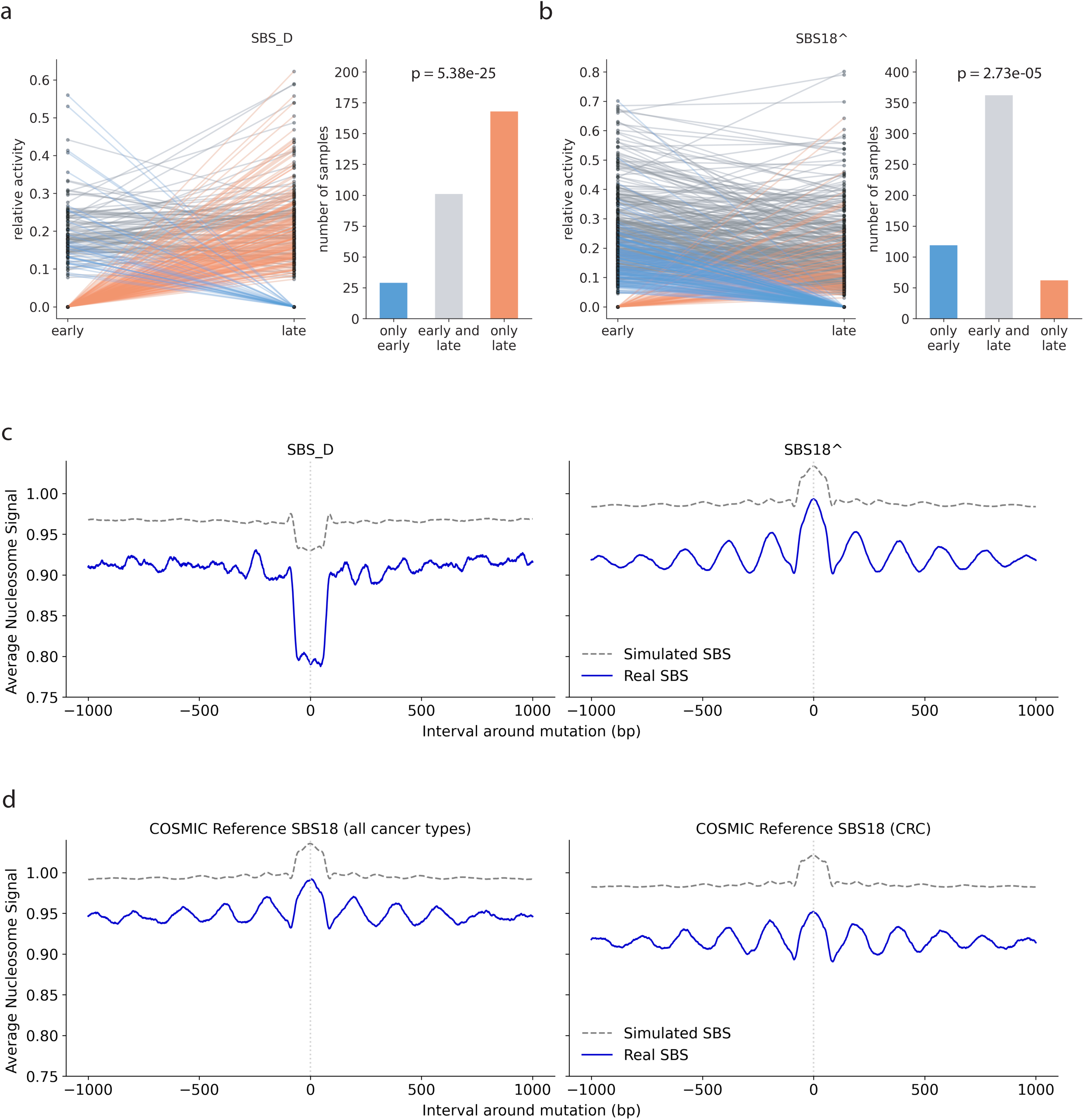
SBS_D and SBS18^ reflect distinct mutational processes. ***(a)*** *(Left)* Relative SBS_D activity in early and late clonal mutations. Each line represents one sample: gray lines indicate SBS_D activity in both early and late clonal mutations, blue lines indicate activity only in early clonal mutations, and red lines indicate activity only in late clonal mutations. *(Right)* Number of samples with SBS_D activity in each category. Samples lacking SBS_D activity in both early and late clonal mutations are excluded from the plots. P-value is based on a McNemar test. ***(b)*** Same format as panel *(a)*, but for SBS18^, showing a different pattern of activity across early and late clonal mutations. ***(c)*** Nucleosome occupancy pattern for SBS_D (*left*) and SBS18^ (*right*), as identified by SigProfilerTopography. Each plot is centered on the mutation site, with 1,000 base pairs upstream (5’) and downstream (3’). Solid lines represent the average nucleosome signal for observed mutations; dashed lines show the signal for simulated mutations. ***(d)*** Nucleosome occupancy patterns for COSMIC signature SBS18 across all cancer types (*left*) and restricted to colorectal cancer (CRC) samples (*right*) from the COSMIC website. Plot features and layout match those in panel *(c)*.

**Figure 3:**
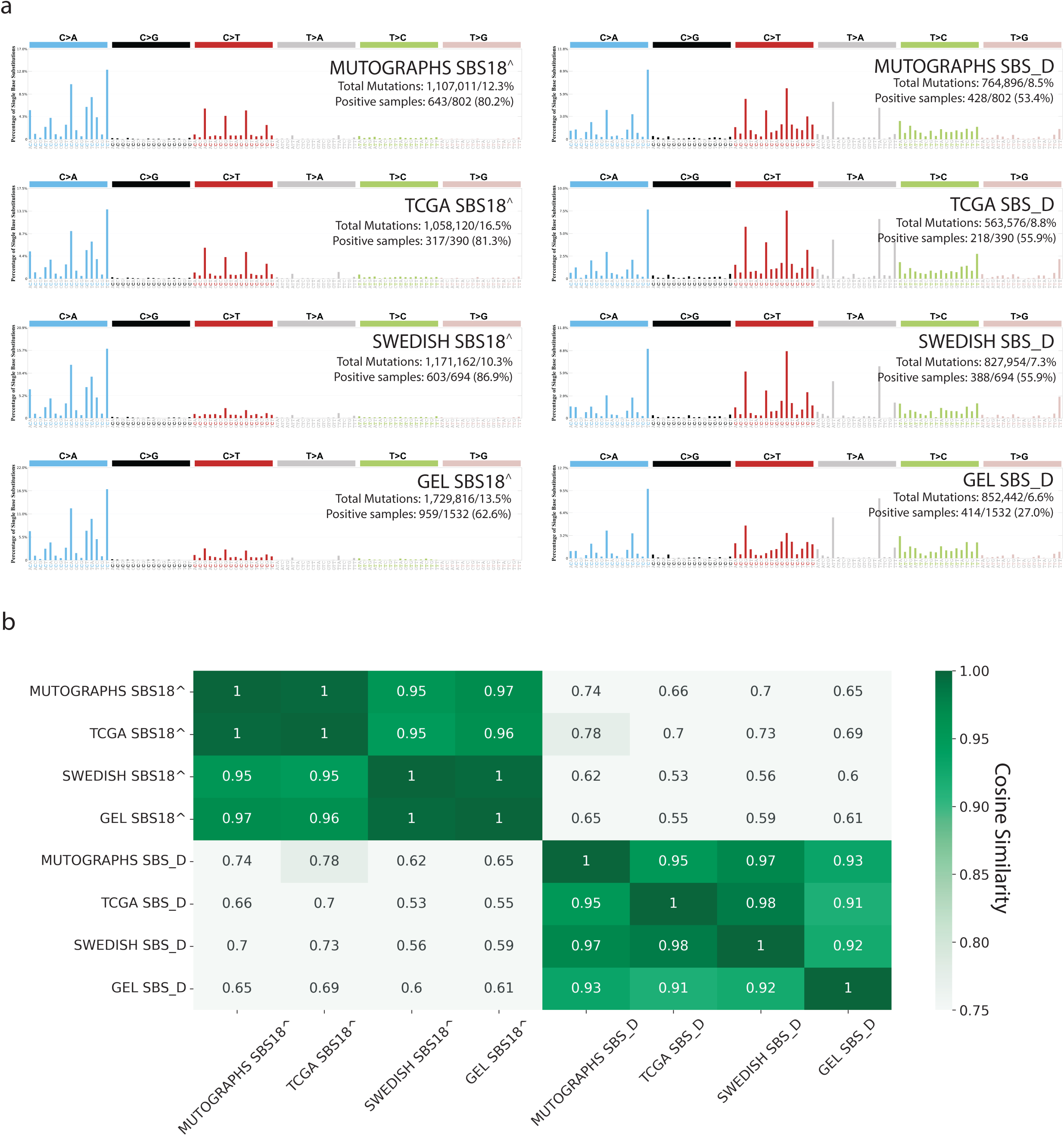
Independent colorectal cancer cohorts support reproducibility of SBS_D and SBS18^. ***(a)*** SBS18^ (*left*) and SBS_D (*right*) signatures extracted from the Mutographs, The Cancer Genome Atlas (TCGA), Swedish, and Genomics England (GEL) colorectal cancer (CRC) cohorts. Signature extraction for the Mutographs, TCGA, and Swedish cohorts was performed using the SBS-288 context and then collapsed to SBS-96 for visualization; the GEL cohort was directly analyzed in the SBS-96 context. The number of mutations and positive samples reflects results from mutational signature attribution (MSA) using *de novo* signatures identified by SigProfilerExtractor. ***(b)*** Heatmap showing cosine similarities among SBS18^-like and SBS_D-like signatures extracted from the four CRC cohorts.

**Figure 4:**
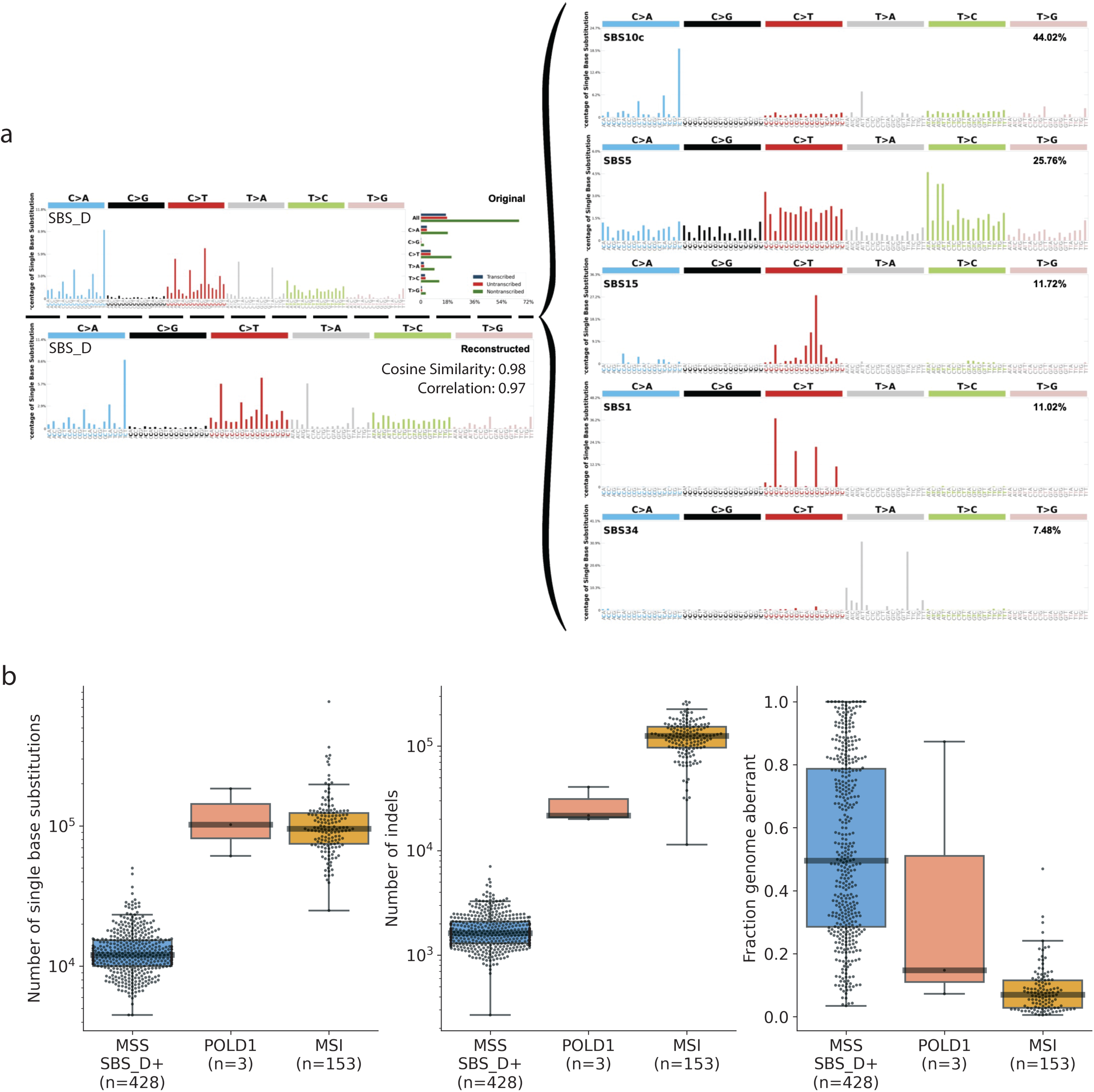
Naïve decomposition of SBS_D and its distinction from *POLD1*-mutant and MSI tumors. ***(a)*** Naïve decomposition of SBS_D using the complete COSMIC reference signature set. ***(b)*** Comparison of microsatellite-stable samples with SBS_D activity (MSS SBS_D+), *POLD1*-mutant samples, and samples harboring microsatellite instability (MSI) in terms of total number of single base substitutions (SBSs; *left*), small insertions and deletions (indels; *middle*), and fraction genome aberrant (*right*).

**Figure 5:**
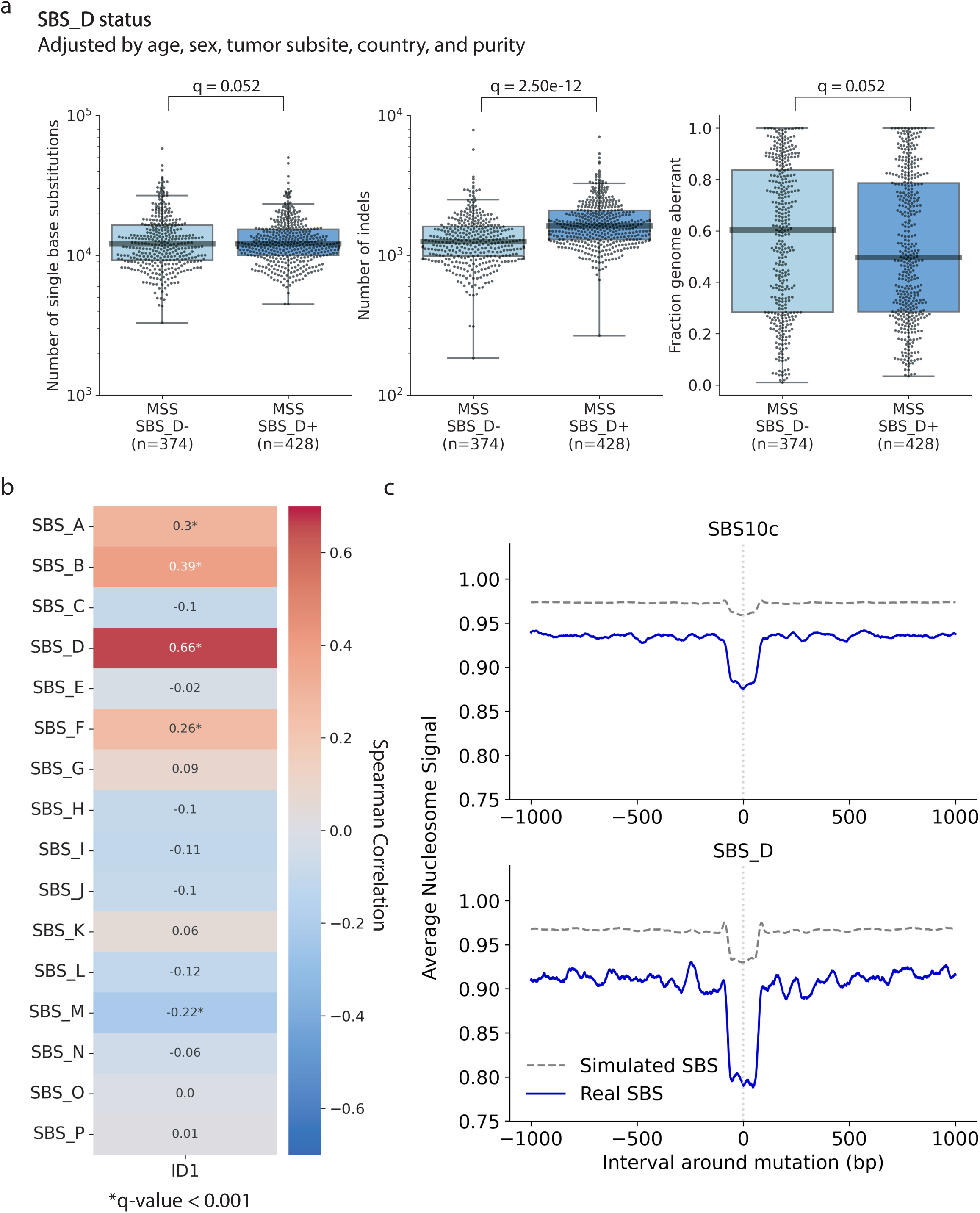
SBS_D exhibits mutational features suggestive of DNA repair infidelity. ***(a)*** Comparison of SBS_D-positive (SBS_D⁺) and SBS_D-negative (SBS_D⁻) samples across total number of single base substitution mutations (*left*), total number of indel mutations (*middle*), and fraction of the genome with copy number aberrations (*right*). P-values were calculated using linear models adjusted for age, sex, tumor subsite, country, and tumor purity, followed by Benjamini-Hochberg multiple testing correction to yield q-values. ***(b)*** Spearman correlation between the activity of each *de novo* SBS signature and ID1 activity. Asterisks indicate significant correlations with q-value<0.001 after Benjamini-Hochberg multiple testing correction. ***(c)*** Nucleosome occupancy pattern for SBS10c in the three *POLD1*-mutant samples from the Mutographs colorectal cancer cohort (*top*); nucleosome occupancy for SBS_D in the microsatellite-stable Mutographs colorectal cancers. Each plot is centered on the mutation site, with 1,000 base pairs upstream (5’) and downstream (3’). Solid lines represent the average nucleosome signal for observed mutations; dashed lines show the signal for simulated mutations.

Additionally, we performed topography analysis on the Mutographs CRC data to elucidate the topographical characteristics of SBS_D and SBS18^ (**Methods**). Although the two mutational signatures were similar in replication timing and strand asymmetry analyses (**Supplementary** Figure 2), they displayed a notable difference in nucleosome occupancy (**Figure 2*c***). Specifically, SBS_D mutations were depleted near nucleosomes and did not exhibit any periodicity. In contrast, SBS18^ mutations were enriched near nucleosomes and demonstrated clear periodicity as expected and previously reported^21^ for COSMIC signature SBS18 in CRC and across all cancer types (**Figure 2*d***). These observations further highlight the clear similarity between SBS18^ and SBS18, and they suggest that SBS_D represents a mutational process dissimilar to the one captured by SBS18.

### Signatures SBS_D and SBS18^ are Validated in Independent Cohorts

Although several large mutational signature extraction studies in CRC have been recently conducted^14,15^, none have reported a *de novo* signature resembling SBS_D. These prior studies analyzed DNA repair–proficient and repair–deficient samples together, which may have masked the presence of an SBS_D-like signature. Consistent with this, when we performed signature extraction on all Mutographs CRC samples without stratifying by DNA-repair status (*n*=981), no signature bearing resemblance to SBS_D was observed (CS<0.85 across all 27 *de novo* signatures; **Supplementary Figure 3**). To validate SBS_D and confirm its distinction from SBS18^, we performed *de novo* signature extraction on treatment-naïve, microsatellite-stable, DNA repair–proficient samples from three independent whole-genome sequenced CRC cohorts, including: *(i)* 390 recently whole-genome re-sequenced samples from The Cancer Genome Atlas (TCGA) project, representing patients predominantly from the United States^13^; *(ii)* 694 samples from a recently published study^14^, representing patients from Sweden; and *(iii)* 1,532 samples from the Genomics England (GEL) project^15^, representing patients from the United Kingdom (**Methods**). *De novo* extractions for the Swedish and TCGA cohorts were conducted using the SBS-288 classification schema, consistent with the approach used in the Mutographs CRC project^9^. In contrast, the GEL cohort was analyzed using the SBS-96 schema, as it is the only publicly available mutational context for these cancers^22^. Despite differences in classification schemas, SBS_D and SBS18^ were identified across all three cohorts with high cosine similarity to their original Mutographs CRC signatures, reinforcing their distinction and confirming the reproducibility of SBS_D extraction in independent DNA repair proficient cohorts (**Figure 3*a-b***). Moreover, aside from the GEL cohort—which showed lower signature prevalence—the total number of mutations and the proportion of samples attributed to these two signatures were very similar across the cohorts, further supporting their robustness and consistency (**Figure 3*a***).

### Naïve Decomposition of SBS_D

The Mutographs CRC study utilized an optimized decomposition method when decomposing *de novo* signatures to COSMICv3.4 signatures^9^. Specifically, since the analyzed samples were treatment-naïve, microsatellite-stable, and DNA repair-proficient, the decomposition process was designed to exclude mutational signatures associated with chemotherapy exposure or DNA repair deficiencies. In the naïve decomposition approach, which allows the use of all known COSMIC signatures, SBS_D decomposed to a mixture of SBS10c (44%), SBS15 (12%), SBS34 (7%), and the clock-like signatures (37%) with cosine similarity of 0.98 (**Figure 4*a***). SBS10c has been shown to strictly come from samples harboring germline or somatic *POLD1* exonuclease domain mutations^23^, and SBS15 has been attributed to defective DNA mismatch repair in microsatellite-unstable samples^12^. At first glance, these results suggest that SBS_D is associated with POLD1 deficiency or microsatellite instability. However, there are at least three lines of evidence that argue strongly against this result. *First*, all *POLD1*-mutant and microsatellite-unstable CRC samples—confirmed by droplet digital PCR—were excluded from the 802-sample cohort prior to signature extraction. We also thoroughly reviewed all samples for any previously missed pathogenic or potentially pathogenic mutations in *POLD1* or the mismatch repair-associated genes. Given that SBS10c has only been observed in tumors with pathogenic germline or somatic mutations in *POLD1*’s exonuclease domain^23^, the absence of such mutations in the 802 samples argues against POLD1 deficiency as the cause of SBS_D (**Supplementary Table 1–2**). Although 12 samples carried non-pathogenic *POLD1* mutations, none showed elevated SBS_D activity (**Supplementary Figure 4a**). Similarly, among the 38 samples with mutations in known mismatch repair genes, SBS_D activity was not elevated (**Supplementary Figure 4b**; **Supplementary Table 1–2**). Only one CRC sample was found to carry a pathogenic *MSH6* nonsense mutation, but it lacked both SBS_D activity and a mutational profile consistent with microsatellite instability (**Supplementary Figure 5**). Similarly, we observed no difference in SBS_D activity between samples with and without loss of heterozygosity in *POLD1* or DNA mismatch repair genes (*POLD1*: 103/774; DNA mismatch repair: 420/774; **Supplementary Figure 4c-d**)*. Second*, while samples with known *POLD1* mutations or microsatellite instability showed 10- to 100-fold higher substitution and indel rates, this elevated mutational burden was not observed in microsatellite-stable samples positive for SBS_D (**Figure 4*b***). *Third*, tumors with pathogenic *POLD1* mutations or microsatellite instability typically have stable, diploid genomes with aberrations affecting less than 20% of the genome, which is in stark contrast to the substantial genomic instability observed in most tumors with SBS_D activity (**Figure 4*b***). While microsatellite instability—but not POLD1 deficiency—can also arise from epigenetic events, mainly the hypermethylation of the *MLH1* promoter^24^, such data were not available for the Mutographs cohort. However, epigenetically driven cases also exhibit high levels of substitutions and indels^25^, which were not observed in SBS_D-positive samples. Together, this supports the conclusion that SBS_D is not driven by canonical defects in POLD1 or mismatch repair.

### Exploring the Potential Etiology of SBS_D

Although SBS_D does not appear to result from canonical defects in *POLD1* or mismatch repair genes, the similarity in mutational patterns suggests that POLD1—and possibly mismatch repair—may still contribute to its underlying mutational processes. This hypothesis is further supported by three key observations in the Mutographs CRC cohort. *First*, although SBS_D-positive samples lacked the characteristic features of POLD1 or DNA repair deficiency—such as hypermutation and low genome aberration (**Figure 4*b***)—they exhibited strikingly higher indel burden compared to SBS_D-negative samples (1.3-fold increase; q-value of 2.50 × 10^-12^ after adjusting for known covariates), whereas negligible differences were observed for SBS burden and genome aberration (**Figure 5*a***). A markedly higher indel burden was also observed in SBS_D positive samples across the TCGA, Swedish, and GEL cohorts (q-values<10^-10^; 1.3-fold increase in TCGA; 1.3-fold increase in the Swedish cohort; 1.4-fold increase in GEL; **Supplementary Figure 6**). *Second*, a prior study^23^ showed that POLD1-deficient tumors are hypermutated specifically for indel signature ID1, which arises from replication slippage of the newly synthesized strand. Notably, SBS_D was the SBS signature most strongly correlated with ID1 activity (CORR=0.66; q-value<0.0001; **Figure 5*b***). *Third*, SBS10c in POLD1-defective tumors exhibits a distinct nucleosome occupancy pattern, marked by strong depletion at nucleosome-bound regions, and a similar depletion pattern was observed for SBS_D in the DNA repair proficient samples (**Figure 5*c***). While we can be confident that SBS_D is not driven by established genomic aberrations in *POLD1* and mismatch repair genes, our results suggest that there may be a link between SBS_D and infidelity in these DNA repair processes.

## DISCUSSION

In this study, we revisited the Mutographs CRC mutational signature analysis^9^, where mutational signature SBS_D was previously decomposed into the reactive oxygen species-associated signature SBS18. Although our results do not alter the main finding from the Mutographs CRC project, they do indicate that the conservative approach taken by Mutographs has missed that SBS_D represents a distinct mutational process from SBS18. Furthermore, although SBS_D is consistently present at similar prevalence across multiple independent microsatellite-stable CRC cohorts, it was not previously reported in these studies. As multiple large-scale colorectal cancer cohorts^14,15^ have previously undergone mutational signature analysis, it is surprising that SBS_D has not been reported. This seems to be due to previous studies not separating DNA repair deficient and DNA repair proficient cancers prior to performing *de novo* signature extraction. DNA repair deficient tumors typically harbor at least an order of magnitude more single base substitutions—around 10^5^ compared to ∼10^4^ in microsatellite-stable cases—which can obscure weaker signals that overlap with signatures of DNA repair deficiency. Since SBS_D shares features with signatures commonly found in hypermutated, DNA repair–deficient tumors, it is particularly prone to being masked during extraction from mixed cohorts. This challenge is further amplified by the fact that SBS_D typically accounts for only 5 to 10% of all mutations in microsatellite-stable tumors. Indeed, without stratifying DNA repair–proficient and DNA repair–deficient samples, the orders-of-magnitude-higher mutation rates from hypermutated microsatellite-unstable or polymerase proofreading-deficient tumors can mask the more subtle contribution of SBS_D. To address this, the original Mutographs CRC study^9^ implemented a key strategy by analyzing microsatellite-stable and unstable samples separately. However, the study also adopted a conservative stance and did not classify SBS_D as a novel signature. At the time, its decomposition into known reference signatures was borderline, and without the additional evidence presented in this study, SBS_D was interpreted as a composite rather than a distinct entity. The results in this study underscore the need to further investigate signatures with borderline decompositions, including analyses of additional cohorts and evaluation of naïve signature decomposition approaches that incorporate all known reference signatures. Overall, despite extensive studies on CRC mutational landscapes, our findings reveal a previously unrecognized mutational signature in microsatellite-stable CRC that may reflect DNA repair infidelity during the late stages of CRC evolution. However, to conclusively determine the etiology of SBS_D, further *in vitro* and *in vivo* experiments will be required. Notably, recent advances in polymerase error rate sequencing (PER-seq) have provided new insights into polymerase epsilon fidelity^26^, and similar approaches will be needed to experimentally validate the potential role of POLD1 infidelity in this newly identified mutational signature.

## METHODS

### Mutational Contexts

Briefly, the SBS-96 mutational context organizes somatic mutations into six base substitution types: C>A, C>G, C>T, T>A, T>C, and T>G (according to the pyrimidine base of the Watson-Crick base-pair). Each type is divided into 16 subtypes based on the trinucleotide context, which accounts for the bases immediately preceding and following the mutation (4 possible bases at the 5′ position × 4 at the 3′ position × 6 substitution types = 96 categories)^17^. The SBS-1536 classification extends the SBS-96 schema by incorporating the pentanucleotide context, resulting in 1,536 mutational channels (256 per each of the six base substitution types). While its complexity has limited its widespread use, SBS-1536 has previously provided insights into mutational processes such as APOBEC-mediated cytidine deamination^27^. The SBS-288 classification builds upon SBS-96 by introducing three additional subcategories for each mutation: *(i)* non-transcribed, (*ii)* transcribed, and *(iii)* un-transcribed. This schema differentiates mutations occurring in non-transcribed (*i.e.*, non-genic) regions of the genome from those in transcribed regions of the genome. Mutations within transcribed regions are further divided based on the transcriptional strand orientation relative to the pyrimidine mutated base^17^, resulting in the *transcribed* and *un-transcribed* classifications. This enhanced framework enables more detailed investigations of transcription-coupled DNA damage and repair processes, for example, transcription-coupled nucleotide excision repair. Importantly, both the SBS-288 and SBS-1536 schemas are fully collapsible to SBS-96, allowing extracted signatures to be converted back to the SBS-96 schema for direct comparison with COSMIC reference signatures. The mutational context enables the transformation of mutational catalogs from cancer genomes into a mutational matrix, where each row represents a specific mutation type and each column corresponds to a tumor sample.

### Mutational Signature Extraction and Decomposition

Mutational signature extraction and decomposition results from the Mutographs CRC Project^9^ were utilized in this study. Briefly, we previously used SigProfilerExtractor^8^, which involves performing nonnegative matrix factorization (NMF) on a mutational matrix, to identify the underlying mutational signatures in the cohort. Extractions were performed using the SBS-96, SBS-288, and SBS-1536 contexts, and the signatures from the SBS-288 extraction were decomposed and used for the Mutographs CRC study. Decomposition of the *de novo* signatures, which means identifying which known reference signatures best explain each of the *de novo* signatures, was performed using an optimized approach where the following signature subsets were removed from the decomposition: artifact signatures, ultraviolet signatures, lymphoid signatures, mismatch repair deficiency signatures, polymerase deficiency signatures, base excision repair deficiency signatures, and treatment signatures. This was done using SigProfilerAssignment^7^ and utilizing the exclude_signature_subgroups parameter.

### Identification of Mutational Signature Activity in Each Sample

As in the original Mutographs CRC project^9^, Mutational Signature Attribution^28^ (MSA) was used to attribute each of the *de novo* mutational signatures to each sample. MSA was run using the same parameters used for the original Mutographs CRC study, including setting the no_CI_for_penalties parameter to false for penalty selection. Analyses were performed using the pruned attributions produced by the tool. Somatic mutations attributed to each mutational signature after mapping to COSMICv3.4—along with SBS_D, which is now classified as a novel signature—are provided in **Supplementary Table 3**.

### Molecular Timing of SBS18^ and SBS_D

Copy number profiles, generated using Battenberg^29^, and somatic variant calls, generated using a consensus between the CaVEMan^30^ and Strelka2^31^ callers, were generated as part of the Mutographs CRC project. These data were used to run DPClust3p^29^ and DPClust^29^ to identify the cancer cell fraction (CCF) of each somatic mutation and estimated number of chromosomes carrying each somatic mutation. The following logic was applied: in areas with copy number amplifications, if a mutation appeared on more than one chromosome copy, it was labeled as early clonal, and if a mutation appeared on one or less chromosome copies when a gain had occurred on the major allele in that location, it was labeled as likely late clonal; this logic was only applied to clonal mutations (those with CCF>0.95). Samples with at least 256 early clonal and 256 late clonal mutations were retained for analysis (684/774), as has been done in previous studies^10,11^. MSA was then used to identify the activity of each *de novo* mutational signature in each early clonal and late clonal set of mutations. A McNemar test was used to assess if the frequencies of samples with SBS_D activity and SBS18^ activity were different between late clonal and early clonal mutations.

### Topography Analysis

SigProfilerTopography^32^ was used to analyze the topographical features of SBS_D, SBS18^, and SBS10c. Probability files and activity files were generated using MSA as part of the Mutographs CRC project. For the SBS_D and SBS18^ analysis, MSA was run with the SBS-288 *de novo* signatures and the 802 DNA repair proficient samples. For the SBS10c analysis, MSA was run using the COSMICv3.4 reference signatures and the three *POLD1*-mutant samples from the full Mutographs CRC cohort. Both runs of SigProfilerTopography^32^ used 0.75 for the average_probability parameter. For replication timing and replication strand asymmetry analyses, the HCT116 colon carcinoma cell line was used as the reference. For nucleosome occupancy, the K562 cell line was used as the reference.

### Somatic Variant Calling in Three Independent Cohorts

For the Swedish Cohort, somatic mutation calls were publicly available and had been identified using Mutect^33^, Mutect2^34^, and TNscope^35^. For GEL, only the SBS-96 mutational matrix was available; somatic mutations for GEL were identified using Strelka^31^. Because mutation calls are not yet available for the TCGA whole-genome sequenced CRC samples, we identified mutations using Mutect2^34^ with the pcr-indel-model parameter set to HOSTILE and all other parameters set to default values; the standard PASS filter was the applied.

### Sample Filtering in Three Independent Cohorts

Samples in each independent cohort were filtered to mimic the DNA repair proficient sample set used in the Mutographs CRC project. For all three cohorts, samples with over 100,000 single base substitution (SBS) mutations or over 7,000 indel mutations were removed. In addition, any samples with less than 1,000 SBS mutations were removed. To account for mutations in *MUTYH*, *NTLH1*, and *OGG1*, samples with cosine similarity greater than 0.80 to any of the respective signatures (SBS36 from COSMIC^36^, SBS30 from COSMIC^36^, and SBS108 from SIGNAL^37^) were removed. Like in the Mutographs CRC project, two samples from the Swedish cohort were removed for suspected microsatellite instable activity (high levels of SBS15) despite not being hypermutators.

### Mutational Signature Extraction in Three Independent Cohorts

After filtering, SBS matrices were generated for the TCGA and Swedish cohort using SigProfilerMatrixGenerator^17^. For the GEL cohort, the publicly available SBS-96 mutational matrix was filtered and used for downstream analysis as the variant calling files are not publicly available. SigProfilerExtractor^8^ with default parameters was then used to extract mutational signatures from each cohort, and the suggested solution was used for each extraction. **Characterizing Pathogenicity of Mutations in *POLD1* and DNA Mismatch Repair Genes** Ensembl Variant Effect Predictor (VEP)^38^ was previously used to annotate mutations in the Mutographs CRC cohort. All mutations with the MODIFIER annotation (lowest impact) were filtered out. All protein-altering single base substitution and indel mutations from the 802 DNA repair proficient samples in *POLD1* and DNA mismatch repair genes (*MLH1*, *MLH3*, *MSH2*, *MSH3*, *MSH6*, *PMS1*, and *PMS2*) were examined using CLINVAR^39^. Results were reported in **Supplementary Table 1** (single base substitutions) and **Supplementary Table 2** (indels).

### Linear Models for Comparing Genomic Features

A generalized linear model (Python version 3.9.7; statsmodels version 0.13.2) was used to identify the association between SBS_D status and total number of single base substitutions, total number of indel mutations, and fraction genome aberrant. Fraction genome aberrant values came from the original Mutographs CRC study^9^. A single model was used for each feature, and sex, age, tumor subsite, country, and purity were all included as covariates. This same approach was used to identify associations between SBS_D status and total number of indel mutations in the Swedish and TCGA cohorts^13,14^. No linear modeling was performed for the Genomics England (GEL) cohort^15^ due to the lack of publicly available metadata.

## SUPPLEMENTARY FIGURE LEGENDS

**Supplementary Figure 1: SBS18^ and SBS_D are recapitulated across signature extractions from multiple contexts in the Mutographs colorectal cancer cohort.** *(a)* SBS_D-like (*top*) and SBS18^-like (*bottom*) signatures extracted from the Mutographs colorectal cancer cohort using the SBS-96 context. *(b)* Same as *(a)*, but signatures were extracted using the SBS-1536 context. *(c)* Cosine similarity between the SBS18^-like signatures and SBS_D-like signatures extracted from each of the SBS-96, SBS-288, and SBS-1536 contexts. All signatures were collapsed to the SBS-96 context for comparison.

**Supplementary Figure 2: SBS_D and SBS18^ exhibit similar patterns for some topographical features. *(a)*** Replication timing patterns for SBS_D (*left*) and SBS18^ (*right*). Blue bars represent deciles (10% bins) of the replication timing signal; the dashed line indicates the distribution of simulated somatic mutations. ***(b)*** Strand asymmetry profiles for SBS_D (*top*) and SBS18^ (*bottom*). Circles indicate statistically significant biases (Fisher’s exact test with Benjamini-Hochberg multiple testing correction), with circle color representing the genomic context. Both signatures show enrichment in intergenic regions (gray circles), while no significant differences were observed between leading versus lagging or between transcribed versus un-transcribed strands. Numbers indicate odds ratios for each comparison.

**Supplementary Figure 3: SBS_D is not extracted when DNA repair proficient and deficient samples are combined.** *De novo* mutational signatures extracted from all the Mutographs colorectal cancer samples (*n*=981), without stratifying by DNA repair status. Extraction was performed using the SBS-288 context; for simplicity, signatures are shown here in the SBS-96 context.

**Supplementary Figure 4: SBS_D activity is not associated with mutations in *POLD1* or mismatch repair genes. *(a)*** SBS_D activity in samples with single nucleotide variants (SNVs, *top*) and indel mutations (*bottom*) in *POLD1*. Statistical significance was assessed using the Mann-Whitney U test for mutation categories with ≥5 samples. ***(b)*** As in *(a)*, but for mutations in DNA mismatch repair genes (*MLH1*, *MLH3*, *MSH2*, *MSH3*, *MSH6*, *PMS1* and *PMS2*). ***(c)*** Comparison of SBS_D activity in samples with and without loss of heterozygosity (LOH) in *POLD1* (*left*) and mismatch repair genes (*right*). Significance was assessed using a Mann-Whitney U test. ***(d)*** SBS_D activity stratified by the number of mismatch repair genes with LOH per sample.

**Supplementary Figure 5: Mutational signature activities in a sample with a likely pathogenic *MSH6* mutation. *(a)*** Mutational profile for sample PD54054a, which carries a likely pathogenic mutation in the DNA mismatch repair gene *MSH6. **(b)** De novo* mutational signatures detected in PD54054a. Numbers on the left of each signature plot represent the number of single base mutations (subs) attributed to that signature. Percentages on the right indicate the COSMIC decomposition of the corresponding *de novo* signature.

**Supplementary Figure 6: SBS_D positive samples have elevated numbers of indels in independent colorectal cancer cohorts.** Comparison of the number of indels between SBS_D-positive (SBS_D⁺) and SBS_D-negative (SBS_D⁻) samples in The Cancer Genome Atlas (TCGA; left), Swedish (middle), and Genomics England (GEL; right) cohorts. For the Swedish and TCGA cohorts, p-values were calculated using linear models adjusted for age, sex, tumor subsite, and purity. For the GEL cohort, the p-value was calculated using a Mann-Whitney U test due to lack of available metadata.

## ACKNOWLEDGMENTS

M.K. was supported by the Cancer Biology, Informatics, and Omics Training Grant (T32CA067754) and the Merkin Fellowship through the University of California San Diego. This work was delivered as part of the Mutographs team supported by the Cancer Grand Challenges partnership funded by Cancer Research UK (C98/A24032). M.R.S. and S.M. are also supported by the Wellcome Trust core grant (206194). M.D.-G. was awarded with a fellowship within the “Generación D” initiative, Red.es, Ministerio para la Transformación Digital y de la Función Pública, for talent attraction (C005/24-ED CV1), funded by the European Union NextGenerationEU funds, through PRTR. D.C.W. is co-funded by Cancer Research UK RadNet Manchester (grant no. C1994/A28701), NIHR Manchester BRC (grant no. NIHR203308), The University of Manchester and the Christie NHS Foundation Trust. This work was supported by the US National Institute of Health grants R01ES032547, R01ES036931, R01CA269919, R01CA296974, P01CA281819, and U01CA290479 to L.B.A. as well as by L,B.A.’s Packard Fellowship for Science and Engineering and the UC San Diego Sanford Stem Cell Institute. The computational analyses reported in this manuscript have utilized the Triton Shared Computing Cluster at the San Diego Supercomputer Center of UC San Diego. The funders had no roles in study design, data collection and analysis, decision to publish, or preparation of the manuscript.

## CONTRIBUTIONS

This study was conceived by M.K. and supervised by L.B.A., D.C.W., M.R.S., and P.B. Analysis of data was performed by M.K. and B.O. with guidance from L.B.A., D.C.W., S.M., M.D.-G, A.A., and S.P. The Cancer Genome Atlas (TCGA) data was downloaded and processed by M.K. and A.A. The manuscript was written by M.K. and L.B.A. with contributions from all other authors. All authors read and approved the final manuscript.

