## Supplementary Figures for "Identification and Validation of a Previously Missed Mutational Signature in Colorectal Cancer"

Supplementary Figure 1

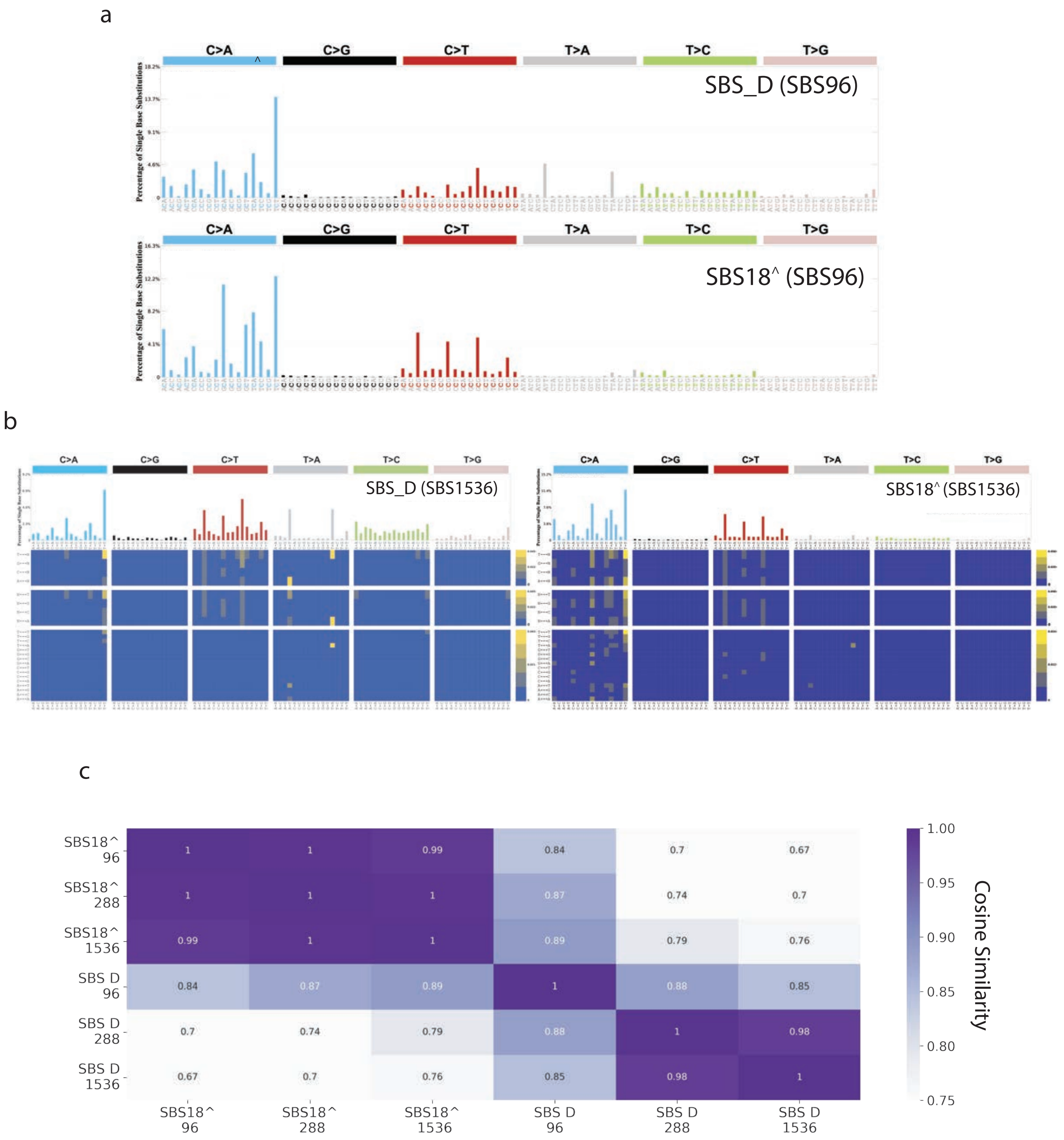

Supplementary Figure 2

a

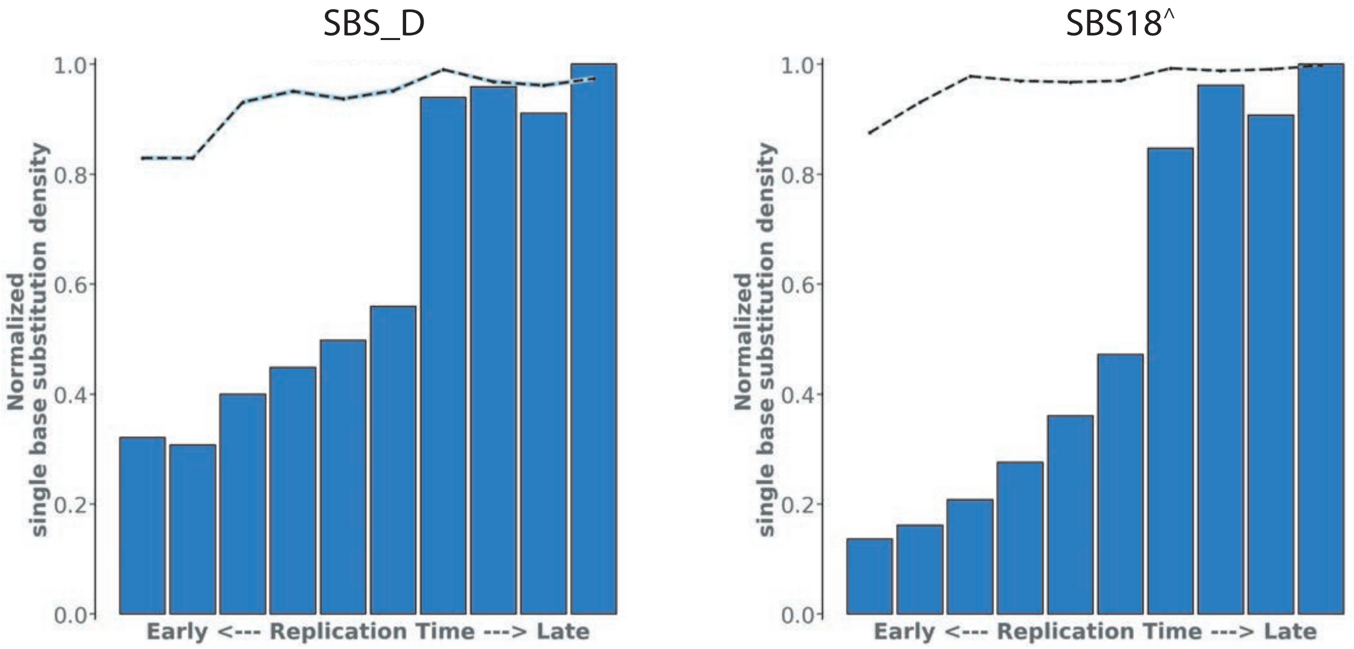


b

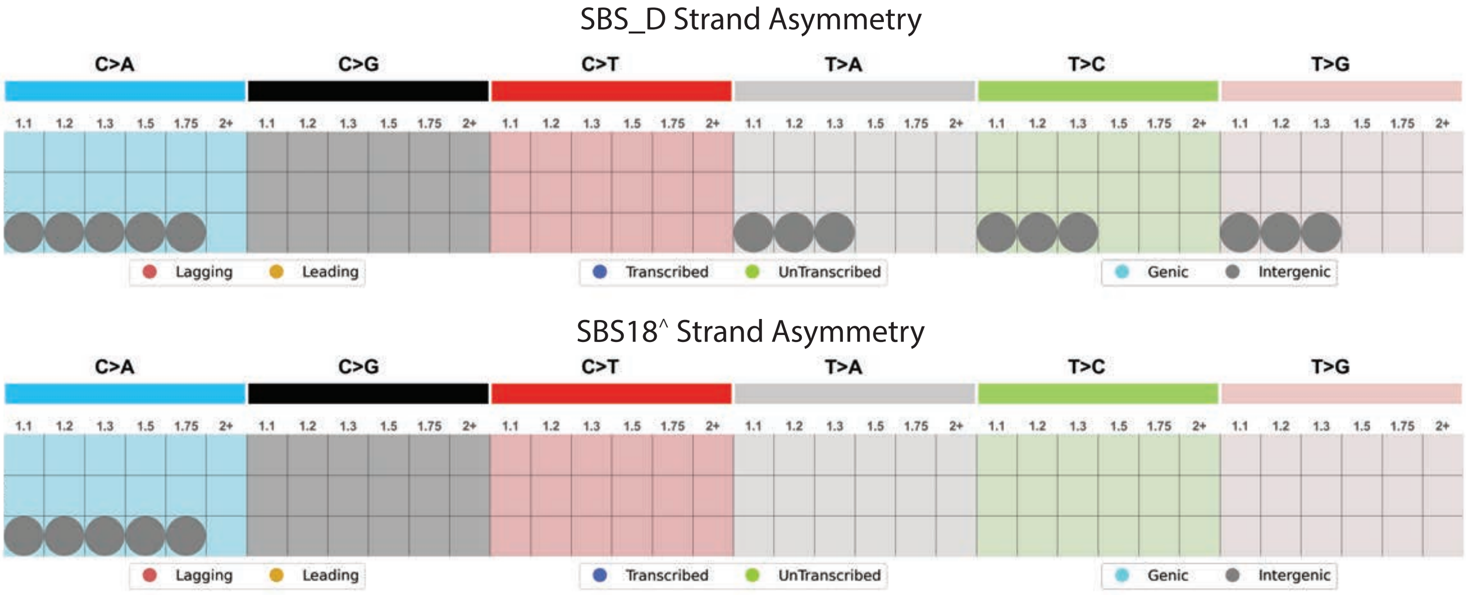

Supplementary Figure 3

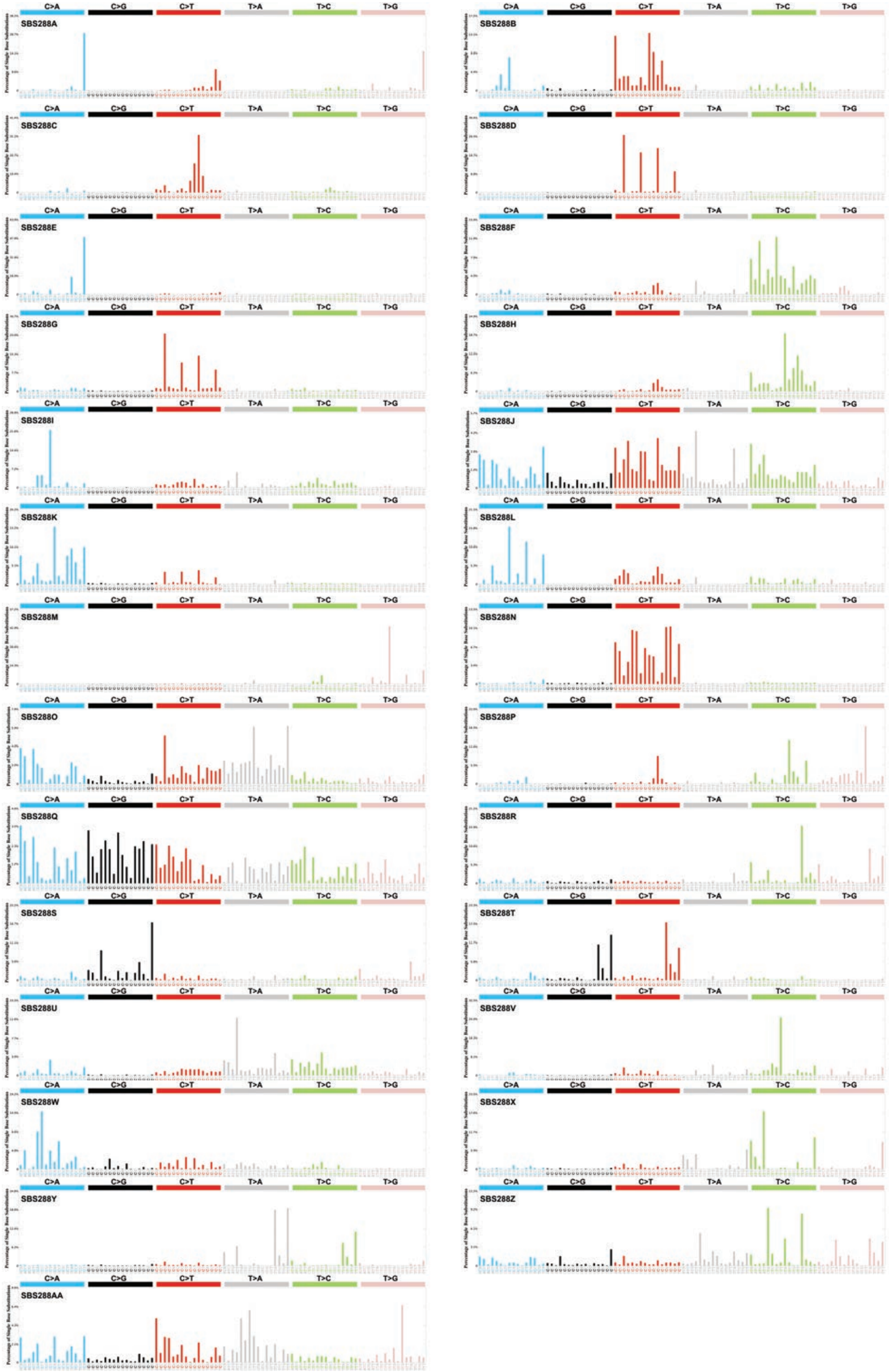

Supplementary Figure 4

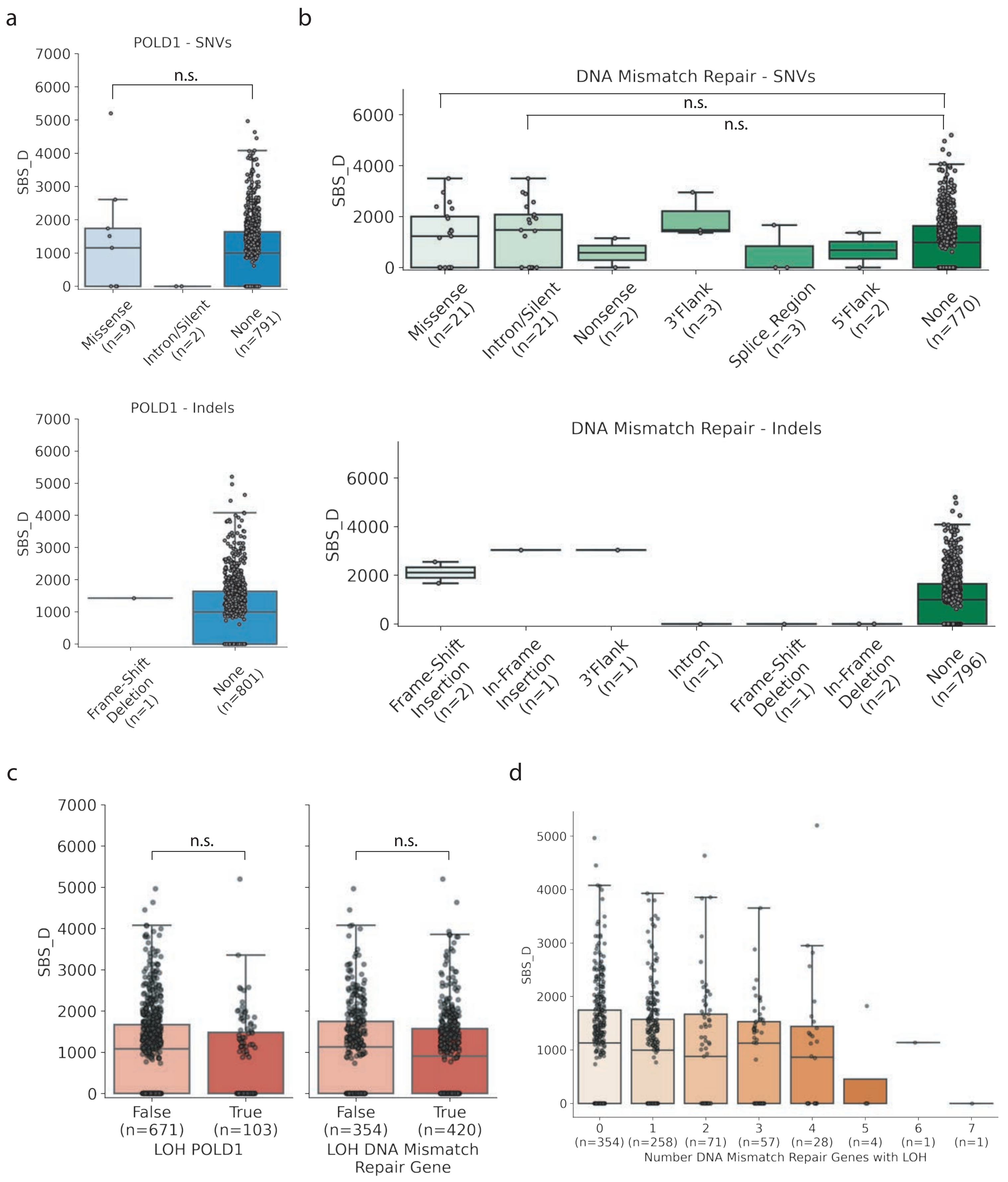

Supplementary Figure 5

a

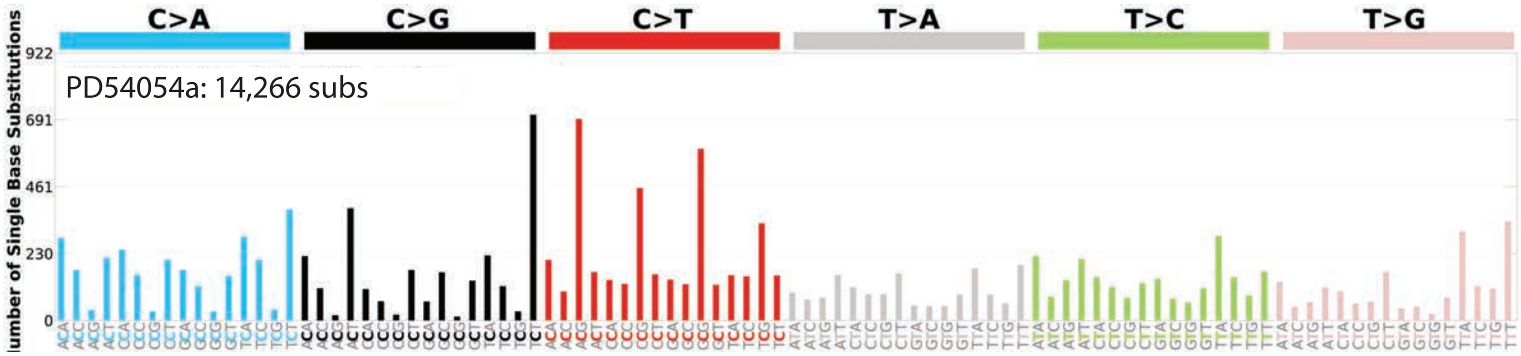


b

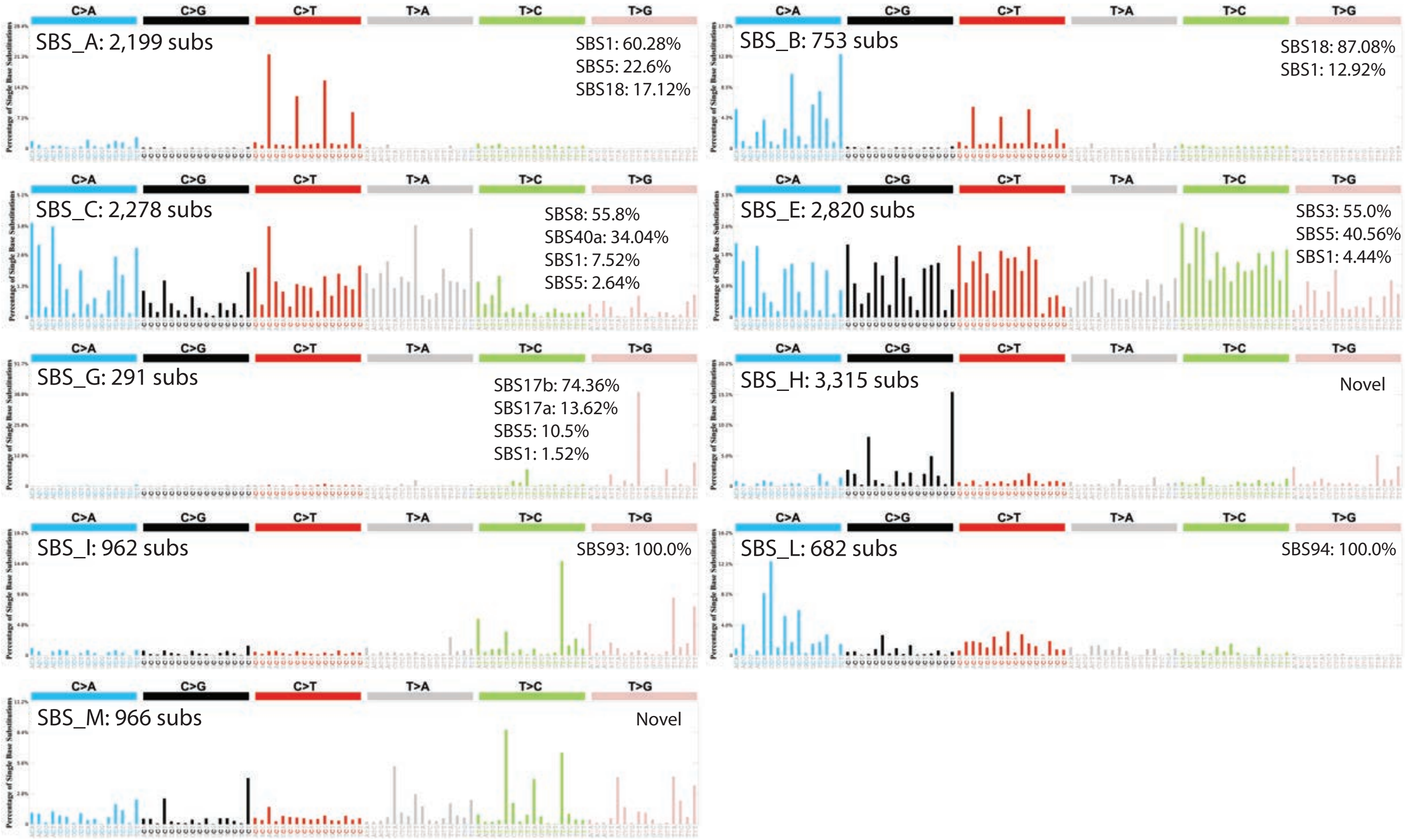

Supplementary Figure 6

**SBS_D status**
Adjusted by age, sex, tumor subsite, and purity (with the exception of GEL)

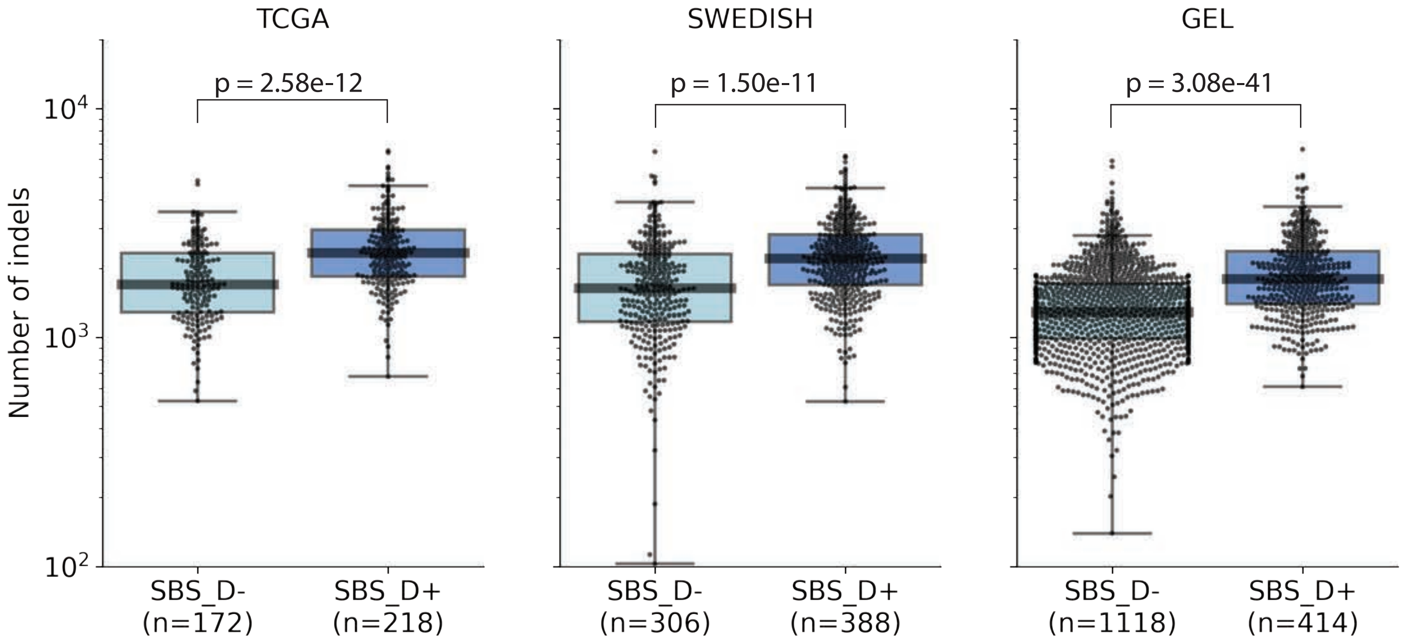
